# Wheat Germ Agglutinin Conjugated Fluorescent pH Sensors for Visualizing Proton Fluxes

**DOI:** 10.1101/781799

**Authors:** Lejie Zhang, Mei Zhang, Karl Bellve, Kevin Fogarty, Maite A. Castro, Sebastian Brauchi, William R. Kobertz

## Abstract

Small molecule fluorescent wheat germ agglutinin (WGA) conjugates are routinely used to demarcate mammalian plasma membranes because they bind to the cell’s glycocalyx. Here we describe the derivatization of WGA with a pH sensitive rhodamine fluorophore (pHRho: pKa = 7) to detect proton channel fluxes and extracellular proton accumulation and depletion from primary cells. We found that WGA-pHRho labeling was uniform, did not appreciably alter the voltage-gating of glycosylated ion channels, and the extracellular changes in pH directly correlated with proton channel activity. Using single plane illumination techniques, WGA-pHRho was used to detect spatiotemporal differences in proton accumulation and depletion over the extracellular surface of cardiomyocytes, astrocytes, and neurons. Because WGA can be derivatized with any small molecule fluorescent ion sensor, WGA conjugates should prove useful to visualize most electrogenic and non-electrogenic events on the extracellular side of the plasma membrane.

## Introduction

Approaches that specifically target fluorescent biosensors to cellular domains are of great interest because they are valuable tools to investigate physiological activity in cells, tissues, and living organisms. Indeed, a review of biosensors that primarily targeted intracellular compartments was published in the JGP in 2017 (Pendin et al., 2017). In contrast, there is a paucity of approaches that target fluorescent biosensors to detect ions and metabolites on the extracellular side of the plasma membrane (Marvin et al., 2013; Patriarchi et al., 2018; Lobas et al., 2019; Sun et al., 2018). A substantial stumbling block is optimizing the protein biosensor to properly fold in the ER and traffic uniformly to the plasma membrane. To avoid the secretory pathway, we previously described a bioorthogonal chemistry approach that targets small molecule fluorescent pH sensors to the cell’s glycocalyx for the real-time visualization of extracellular proton accumulation and depletion (Zhang et al., 2016). Although our first approach detected proton fluxes at the plasma membrane from ion channels and membrane transport proteins, it required cells to utilize and efficiently incorporate an unnatural sialic acid precursor into the glycocalyx. In addition, the calculated pH values were approximations because we had to assume the cell surface labeling was constant over the entire plasma membrane. Therefore, we sought a simplified approach that would work with most cells and provide calibrated changes in extracellular pH adjacent to the plasma membrane.

Small molecule-wheat germ agglutinin (WGA) conjugates appear to be well-suited to target biosensors to the glycocalyx because fluorescent WGA conjugates are commonly used to demarcate the plasma membrane of fixed cells. Although prone to photobleaching and internalization, WGA-fluorescein has been used to approximate extracellular pH at the glycocalyx (Villafuerte et al., 2014; Stock et al., 2007; Schroeder et al., 2013), raising the possibility that pH-sensitive WGA-conjugates may enable the visualization of extracellular proton fluxes from voltage-gated ion channels, membrane transport proteins, and proton omega currents that have been associated with human disease (Fig. 1).

**Figure 1.**
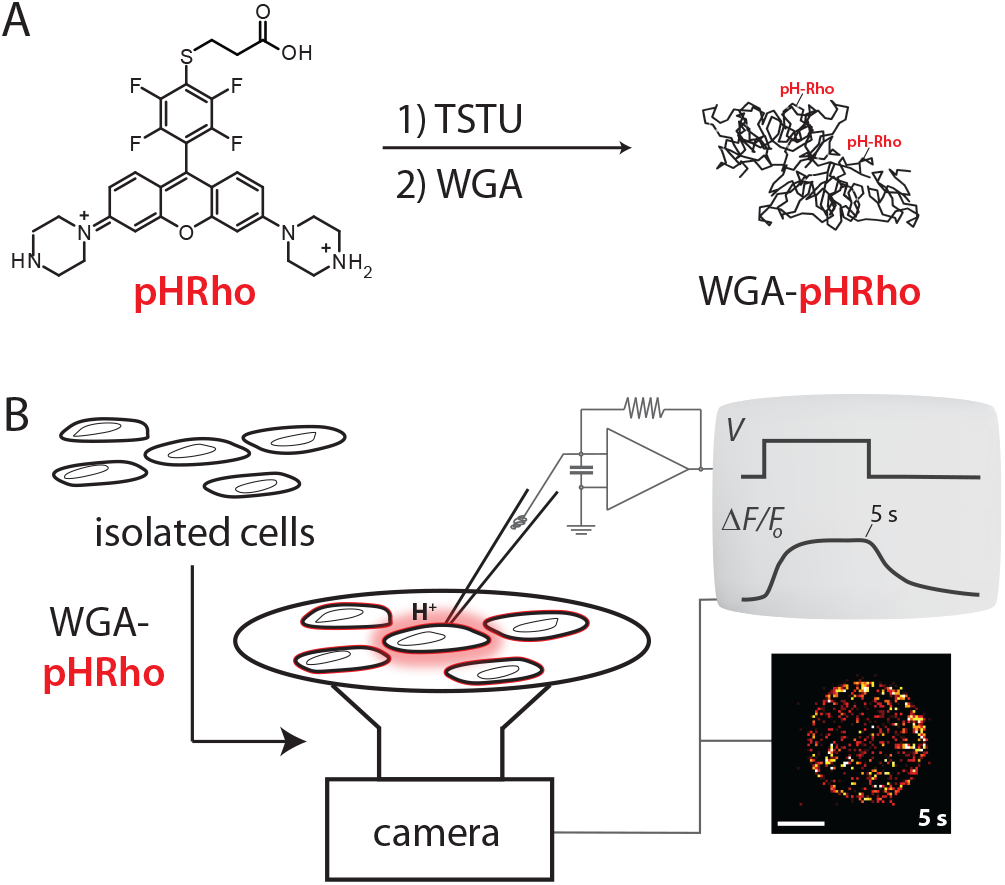
Derivatization of Wheat Germ Agglutinin (WGA) with a small molecule fluorescent pH sensor and cell surface labeling to visualize plasma membrane proton fluxes: (**A**) Synthetic scheme of WGA-pHRho. (B) Cartoon depiction of the whole cell patch clamp fluorometry approach to visualize proton fluxes with WGA-pHRho.

Here we describe the synthesis and utilization of WGA-pHRho (Fig. 1A), a pH-sensitive, red fluorescent WGA conjugate to visualize extracellular proton accumulation and depletion. Although WGA binds to the terminal sugars of glycosylated proteins, labeling cells with WGA conjugates did not appreciably alter the voltage-gating of glycosylated or non-glycosylated ion channels. Extracellular proton fluxes were fluorescently detected from cells coated with either WGA-pHRho or commercially-available WGA-fluorescein 4 hr after labeling despite visual signs of endocytosis. Compared to WGA-fluorescein, WGA-pHRho was photostable, increased fluorescence upon protonation, and has a pKA ~ 7 that is well-matched for extracellular pH determinations. On-cell calibration of WGA-pHRho and WGA-fluorescein provided accurate changes of pH over the extracellular surface of mammalian cells. Using various imaging modalities, differential extracellular proton accumulation and depletion was detected at (i) the sarcolemma and t-tubules of cardiomyocytes and (ii) in neuron-astrocyte co-cultures. This straightforward approach and the detailed procedures for WGA conjugation and cell surface labeling will enable physiologists and biophysicists to employ these reagents to fluorescently visualize proton fluxes and to spatiotemporally determine extracellular pH changes adjacent to the plasma membrane.

## Materials and Methods

### WGA-pHRho synthesis

pH-Rho was synthesized as described previously (Zhang et al., 2016). pH-Rho (1.0 mg, 1.4 μmol) was reacted with 14 μmol of N,N,N’,N-Tetramethyl-O-(N-succinimidyl) uronium tetrafluoroborate (TSTU) in 200 μL of anhydrous N,N-dimethylformamide (DMF) at RT for 1 hr. The above reaction mixture (100 μL; 5:1 dye:protein molar ratio) was added to 1 mg of WGA (Vector Laboratories) in 1 mL of 0.1 M sodium bicarbonate (pH 8.3), containing 120 mg N-acetylglucosamine. The reaction mixture was stirred at RT for 2 hr and was terminated by adding 6.25 μL of a hydroxylamine solution to a final concentration of 0.1 M. For WGA-fluorescein, a FITC stock solution was prepared at 10 mg/mL in DMF immediately before starting the reaction, and 0.26 μmol FITC dye was added to WGA solution with N-acetylglucosamine (pH 9.0), final dye-to-WGA ratio was 10:1. The WGA-conjugates were separated from unreacted fluorophore by gravity gel filtration on a Sephadex G-25 Fine (20 – 50 μm) column equilibrated in PBS, final samples were lyophilized down to a powder. The degree of labeling (D.O.L.) was calculated using the following equation: the absorbance of WGA-conjugate was measured at 280nm (A_280_) and the λ_max_(A_max_).

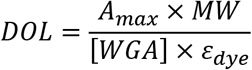

where MW is the molecular weight of WGA, *ε_dye_* is the extinction coefficient of the dye at its absorbance maximum, and the WGA protein concentration is in mg/mL. WGA-fluorescein was purchased from Vector Laboratories.

### CHO cell labeling with pH sensitive WGA conjugates, cell surface calibration, and wholecell patch-clamp fluorometry

CHO cell culture and transfection procedures were conducted as previously described (Zhang et al., 2016). Transfected CHO cells were trypsinized and seeded on a glass-bottomed culture dish for 2 h, labeled with WGA conjugates (50 μg/mL) in HBSS at RT for 10 min, and then rinsed with bath solution (in mM): 145 NaCl, 5.4 KCl, 2 CaCl_2_, and 0.1 buffer (HEPES for pH 7.5, MES for pH 6.0) with NaOH. Whole-cell VCF was performed (Zhang et al., 2016) using internal solution (in mM): 126 KCl, 2 MgSO_4_, 0.5 CaCl_2_, 5 EGTA, 4 K2-ATP, 0.4 guanosine triphosphate, and 25 buffer (HEPES for pH 7.5; MES for pH 6.0). Cells were imaged at 10Hz using a CoolLED pE-4000 light source (exposure time 10ms), 63 × 1.4 numerical aperture oil-immersion objective, and a Zyla sCMOS camera (ANDOR). WGA-fluorescein and WGA-pHRho were excited respectively at 490 and 550 nm channels (Chroma 89402 emission filter set). Fluorescent images were collected (4×4 binning) and processed using ImageJ software.

### Rat ventricular myocyte isolation, WGA-pHRho labeling, and structured light microscopy

Rat ventricular myocytes were isolated from adult Sprague-Dawley rats (200 – 250 g) by enzymatic (Liberase™, Roche) digestion (Colecraft et al., 2002). Animals were anaesthetized with *Katamine/Xylazine* under the guidelines of UMASS Medical School Animal Care and Use Committee. Hearts were excised and ventricular myocytes were isolated using Langedorff perfusion. Healthy, rod-shaped cardiomyocytes were cultured in Medium 199 on laminin-coated glass bottom culture dishes with 5% CO_2_ and 95% air at 37 °C for 2 h before imaging experiments. Cardiomyocytes were incubated with WGA-pHRho (50 μg/mL) in HBSS buffer at RT for 30min. Cells were first perfused with Tyrode’s solution (in mM): 145 NaCl, 5.4 KCl, 2 CaCl_2_, 1 MgCl_2_ and 0.1 HEPES, pH 7.5 and then with 10 mM lactate in Tyrode’s. The MCT-1 inhibitor, AR-C155858 (50 nM) was incubated 30 min at RT before lactate perfusion. Cardiomyocytes were imaged on TESM, a custom-built microscope system that has TIRF and structured illumination wide-field epifluorescence for fast optical sectioning and enhanced spatial resolution (Navaroli et al., 2012). The structured light images were formed by illuminating through a grating (500 lines-pairs/in) on a Physik Instruments translation stage that was moved one-third and two-thirds of a period (∼2 ms) for the second and third images. The three structured light images (180 sets of 3 x 50 ms exposures at 1 Hz) were collected at a focal plane 1.8 μm above the coverslip on an inverted Olympus IX71 microscope with a 60 × 1.49 numerical aperture oil-immersion objective with a 1.6X optivar. The sample was illuminated with a Cobolt diode pumped solid state (DPSS) 100mW laser (561 nm) and the emission collected onto a 1004×1002 Andor iXon 885 emCCD camera, which was binned at 2×2.

### Structured illumination image processing and concentric annuli analysis

All image processing was performed using custom software, but the operations are available in ImageJ. At each time point, the three unprocessed structure illumination grating position images were added (corresponding x, y pixels) to reject out-of-focus light (Neil et al., 1997) and produce a wide-field equivalent image with higher signal-to-noise. The processed images were corrected for any remaining locally diffuse fluorescence background by subtracting the morphological opening (maximum of the minimums) of each time point using a radius of 10 pixels (1.67 μm), which removes the local background while preserving bright objects smaller than ~3 μm. To generate a closed (2-D) outline of the cell, a maximum intensity projection was made from the first 30 preperfusion frames of the time series and the outermost visible boundary of the sarcolemma manually trace. The concentric binary (0|1) annular masks were generated by creating a mask of the entire cell and applying a morphological binary erosion of 5 pixels (835 nm), then subtracting this smaller mask from the original to create annular mask A0. Annular masks A1 – A3 were generated by repeating the process using the smaller mask as the starting point. Using these 2-D annular masks, the total and average (normalized for area) fluorescence signal of each annulus was calculated at each time point of the processed image series. The fluorescent signal from the whole cell was calculated using the cell outline mask. The computed average signal over time, F(t) for the whole cell and each annulus, was converted to percent change in signal ΔF/F_0_ = 100*[F(t) – F_0_]/F_0_ using the average of the pre-perfusion time points as F_0_.

### WGA-pHRho labeling and imaging of primary astrocyte and neuron co-cultures

Astrocyte and neuron co-cultures were obtained from P1 pups of C57bl/6 mice. All C57bl/6 (females and males, 1 day old) were obtained from the Biosecurity Laboratory Facility for Animal Experimentation, Janelia Research Campus, HHMI). All procedures were conducted in accordance with protocols approved by the Janelia Institutional Animal Care and Use Committee. Forebrains were removed and the cortex dissected. Tissue was digested with 0.12% trypsin (wt/vol, Gibco Co., Rockville, MD, USA) in 0.1 M phosphate buffer (PBS: pH 7.4, osmolarity 320 mOsm) and mechanically disrupted with a fire-polished Pasteur pipette. Cells were plated at 0.3 x 10^6^ cells/cm^2^ in glass bottom dishes (Cellvis) coated with poly-L-lysine (mol. wt > 350 kDa, Sigma-Aldrich) and cultured for up to 10 days in Nbactive4 (BrainBits®) medium supplemented with 3% fetal bovine serum. Cultured cells were incubated for 30 min in 500 μL of glucose & buffer free media (GBFM) containing (in mM): 140 NaCl, 20 KCl, 2 CaCl_2_, and 2 MgCl_2_. During the last 20 minutes of glucose deprivation, 5 μL of WGA-pHRho (1 mg/mL) was added to the incubation media. The cells were rinsed three times with GBFM and imaged. Imaging was performed at 37 °C, 5% CO_2_ chamber on an inverted Zeiss LSM 880 microscope with a 63 × 1.4 numerical aperture oil-immersion objective equipped with an Airyscan array detector. The astrocyte and neuronal planes were imaged at 2 Hz (50 ms exposures) using 405 and 561 nm laser lines for DAPI and WGA-pHRho, respectively. Glucose (3 mM final concentration) and rotenone (100 μM final concentration) were directly added to the recording chamber on each running experiment. Change in intensity values were calculated in ImageJ using maximum intensity projection series. Glucose and rotenone were obtained from Sigma-Aldrich. Acquisition, visualization and airyscann post processing were performed using Zen software (Zeiss), analysis was performed with ImageJ, and data plots were generated with Microcal Origin 9.

## Results

We were initially concerned that WGA binding to the terminal sugars of a cell’s glycocalyx would drastically alter voltage-gating by either binding to glycosylated voltage sensors or shielding the negatively charged sialic acids that have been implicated in voltage-gating (Johnson and Bennett, 2008; Johnson et al., 2004; Schwetz et al., 2011; Noma et al., 2009). To test the effects of WGA on voltage-gated ion channels, we recorded families of currents from cells transiently transfected with voltage-gated proton (hHv-1) or potassium (Shaker-IR) potassium channels before and after treatment with WGA (Fig. 2A). Hv-1 cannot be N- or O-glycosylated (it does not contain any extracellular threonines or serines) whereas Shaker-IR contains two N-linked glycosylation sites in the S1-S2 loop of the voltage-sensor. Tail current analysis showed that WGA treatment did not significantly affect the midpoint of voltage activation (V_1/2_) of Hv-1 (Fig. 2B). WGA perfusion (Fig. 2A, middle) did slightly alter Hv-1 activation and deactivation kinetics; however, these subtle changes were due to WGA binding to endogenous glycoproteins because Hv-1 does not have any extracellular serine or threonine residues and cannot be glycosylated. In contrast, Shaker-IR activation/deactivation gating was unaffected by WGA perfusion. WGA binding to N-glycosylated Shaker-IR channels did cause a small, but statistically insignificant shift in V_1/2_ (Fig. 2B).

**Figure 2.**
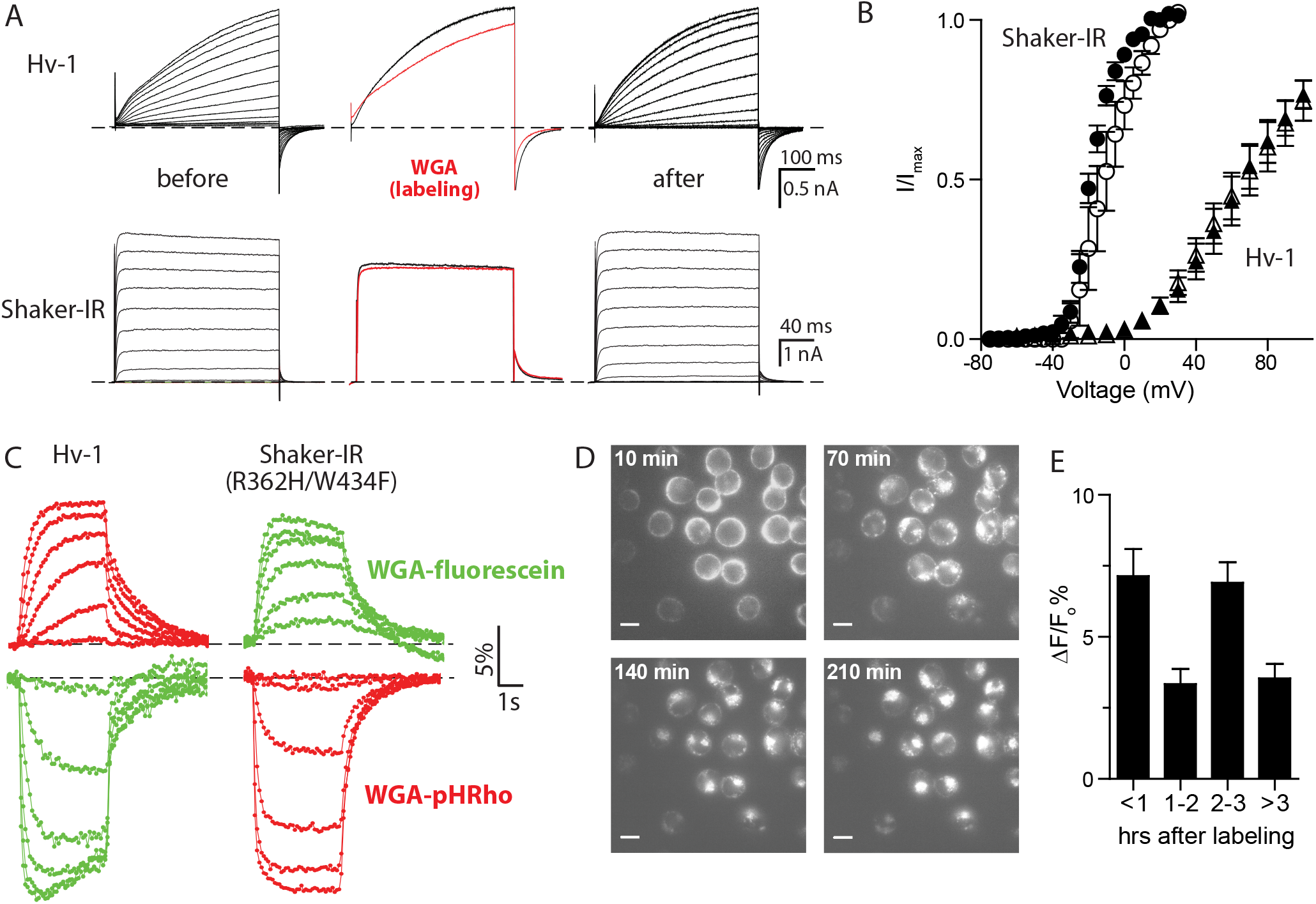
WGA labeling and voltage clamp fluorometry of CHO cells expressing voltage-gated channels. (**A**) Hv-1 (*Top*) and Shaker-IR (*Bottom*) currents were recorded before (*left*) and after (*right*) WGA labeling (50 μg/mL). *Middle* panels are currents before (black) and after (red) WGA wash-in at 60 mV. Cells were held at – 80 mV, Hv-1 depolarized for 0.5 sec from – 50 to 100 in 10-mV increments and Shaker-IR depolarized for 0.2 sec from – 70 to 60 mV in 10-mV increments. (**B**) G-V curves of Hv-1 (V_0.5_ = 68 ± 6 mV, n=8, triangles) and Shaker-IR (V_0.5_= – 18 ± 1 mV, n=5, circles) before (closed) and after (open) WGA treatment. Hv-1: V_0.5_ = 67 ± 8 mV, n=9; Shaker-IR: V_0.5_= – 11 ± 1 mV, n=6. (**C**) Voltage-clamp fluorometry traces from cells labeled with WGA-pHRho (red) and homemade WGA-fluorescein (green). Hv-1 (*left*) was held at – 80 mV and changes in fluorescence were elicited from 4-s depolarizations from 0 to 100 mV in 20-mV increments. Shaker-IR R362H (*right*) was held at + 30mV and fluorescence were elicited from 4-s command voltages from – 120 to – 20mV in 20-mV increments. (**D**) Representative fluorescent images of CHO cells 10 – 210 min after labeling with WGA-pHRho for 30 min; scale bars are 10 μm. (**E**) Bar graph of ΔF/F_0_(%) vs time after WGA-pHRho labeling.

Given that unglycosylated and glycosylated voltage-gated ion channels were minimally affected by WGA treatment, we next determined whether pH-sensitive WGA conjugates would effectively report on proton accumulation and depletion on the extracellular side of the plasma membrane. We made a red fluorescent, pH-sensitive WGA-conjugate (WGA-pHRho) by first synthesizing the NHS-ester of pH-Rho (DCC in DMSO) (Fig. 1A) and then adding the reaction mixture to WGA for 2 h. The reaction was quenched with hydroxylamine and the unreacted dye and excess N-acylgalactosamine (to protect the amino groups in the carbohydrate binding sites) were removed with a gravity-fed size exclusion column (Materials and Methods). Conjugation of the small molecule pH-sensitive fluorophore to WGA did not change the apparent pKa 6.9 or proton-dependent change in fluorescence (Fig. S1A). Based on the protein-fluorophore absorbance ratio, the degree of labeling was 1.0 – 1.1 pHRho molecules per WGA protein.

To fluorescently visualize proton efflux with WGA-pHRho, cells were treated with 50μg/ml of WGA-pHRho for 10 min and the currents and fluorescence were measured using patch-clamp fluorometry in a bath solution with a low buffer capacity (0.1 mM). The top left panel in Figure 2C shows families of fluorescent signals from a CHO cell expressing a C-terminally GFP-tagged, human voltage-gated proton channel (Hv-1) that was labeled with WGA-pHRho. Under these low buffering conditions, voltage-activation of Hv-1 channels resulted in small currents (Fig. S2A) that reached steady state faster with stronger depolarizations.

Simultaneous imaging of the cell’s fluorescence at 10 Hz showed that the change in fluorescence was similar to voltage-dependence of Hv-1; however, the rate to reach steady state was slower as it corresponded to proton accumulation at the cell surface and not Hv-1 channel gating. This distinction between channel gating and proton accumulation (or depletion) was evident upon Hv-1 deactivation at – 80 mV: no current remained after 300 ms whereas it required seconds for the fluorescent signal to fully decay. As we have had previously observed (Zhang et al., 2016), the fluorescent signal decay has two time constants: the fast component corresponds to protons rushing into the cell before Hv-1 channel closing; the slow component corresponds to diffusion into bulk solution. A movie of F/F_o_ snapshots before, during, and after a 100-mV test pulse highlights the kinetic differences in proton accumulation and depletion at the cell surface (Movie S1).

To visualize proton depletion using WGA-pHRho, we set up an inward proton gradient and used a potassium pore-blocked (W434F) Shaker-IR mutant (R362H) that creates an omega proton pore upon hyperpolarization (Fig. 2C, lower right panel). As expected for inward proton flux, the cell surface fluorescence became dimmer upon hyperpolarization, reached steady state, and then recovered with biexponential kinetics when the cell was returned to the 30-mV holding potential (Movie S2). All fluorescence changes were due to proton channel expression (Fig. S2B), as no change in fluorescence was observed when the cells were transiently transfected with empty plasmid DNA (Fig. S2C).

We next compared WGA-pHRho to WGA-fluorescein (Fig. 2C). For a fair comparison, we made our own WGA-fluorescein by reacting commercially available FITC with WGA and removed the unreacted fluorophore with a gravity-fed size exclusion column (Methods and Materials). Compared to WGA-pHRho, homemade WGA-fluorescein (Fig. 2C, green traces) yielded the mirror opposite response to extracellular proton accumulation (Hv-1) and depletion (Shaker-IR R362H/W434F) with approximately the same current-ΔF/Fo relationship (Fig. S2B). As expected, photobleaching was problematic with WGA-fluorescein (the data in Fig. 2C are uncorrected), especially for extracellular proton accumulation where the fluorescent signal was also quenched by protons. There was no noticeable difference between commercially available and homemade WGA-fluorescein (Fig. S2B); thus, for extracellular proton depletion near fluorescein’s pkA of 6.4, commercially-available WGA-fluorescein was a satisfactory reagent when photobleaching can be reliably corrected.

The exemplars in Fig. 2C were performed within 1 hr of labeling with WGA-pHRho and WGA-fluorescein, which was not technically demanding, but we wondered whether internalization of WGA conjugates would affect the fluorescent signals at the plasma membrane. Up to 1hr after labeling, the majority of the WGA-pHRho appeared to be on the cell surface as evidenced by the circular labeling (Fig. 2D); however, bright puncta, indicative of WGA-pHRho internalization became evident 2 hr after labeling, and visually overwhelming at 4 hr. Surprisingly, the change in fluorescence did not systematically diminish over time, indicating that after 4hr there were enough pH sensors at the cell surface to detect extracellular proton accumulation and depletion. Part of the surprise was due to the optical illusion created by the overemphasized endocytosed sensors that were localized in low pH internal compartments (endosomes and lysosomes). In addition, the data in the latter timepoints likely favored cells with less internalized WGA-pHRho because it became noticeably harder over time to make and maintain a gigaohm seal in the whole cell configuration. Thus, it was possible to visualize proton fluxes from WGA-pHRho treated cells 4 hr after labeling; however, for experimental ease, it was experimentally prudent to image and record immediately after labeling.

To convert the change in fluorescence into pH, we perfused pH standards across cells labeled with WGA-pHRho and WGA-fluorescein and plotted ΔF/F_0_ versus pH (Fig. 3A). Perfusion of pH standards with 1 mM or 10 mM buffer yielded linear responses for both fluorescent WGA-conjugates. Using the linear fits, the fluorescent signals for Hv-1 were converted into ΔpH (Fig. 3B). The snapshots at 5 s at 0, 20, 100 mV highlight the advantage of the fluorescence response of WGA-pHRho for detecting extracellular proton accumulation (Fig. 3C). In addition to seeing proton efflux from the patch clamped cell (dotted circle), the acidification of the bath solution was revealed by the neighboring WGA-pHRho labeled cells in Figure 3C.

**Figure 3.**
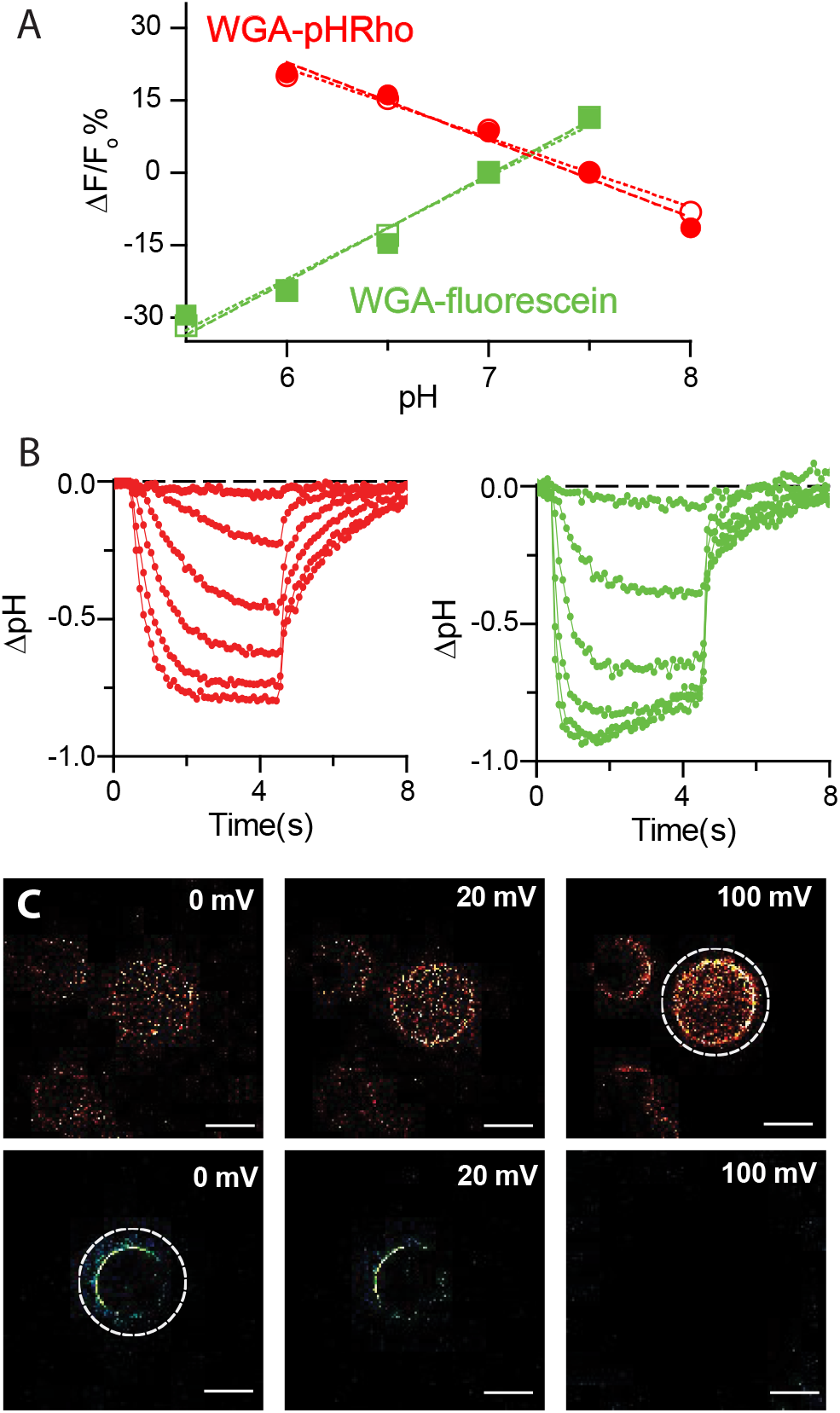
Cell surface pH calibration of WGA-pHRho and WGA-fluorescein. (**A**) CHO cells were labeled with either WGA-fluorescein or WGA-pHRho and the change of fluorescence (ΔF/F_0_%) was plotted using pH standards (HEPES: 1 mM, open symbols; 10 mM closed symbols); Fo was pH 7.5 for WGA-pHRho; pH 7.0 for WGA-fluorescein. The linear fits for WGA-pHRho were: pH = 7.49 – 0.06*[ΔF/F_0_] and pH = 7.41 – 0.07*[ΔF/F_0_] for 1 and 10 mM, respectively; WGA-fluorescein: pH = 7.01 + 0.05*[ΔF/F_0_] and pH = 7.02 + 0.05*[ΔF/F_0_] for 1 and 10 mM, respectively. (**B**) Conversion of ΔF/F_0_ (%) to CHO cells expressing Hv-1 were held at – 80 mV and changes in fluorescence were elicited from 4-s depolarizations from 0 to 100 mV in 20-mV increments and converted into ΔpH using the calibration curves in **A**; WGA-pHRho: red; WGA-fluorescein: green. (**C**) ΔF/F_0_ images of cells expressing Hv-1 taken at 4 s shown in **B**. *Top*: WGA-pHRho; *bottom*: WGA-fluorescein. Dotted white circles indicate the voltageclamped cell.

Because fluorescent WGA conjugates have been used to demarcate the plasma membrane of isolated cardiomyocytes, we next determined whether WGA-pHRho could differentially detect lactate-coupled proton transport in the t-tubules and at the sarcolemma (Fig. 4). Freshly isolated rat ventricular myocytes were labeled with WGA-pHRho for 30 min, rinsed with Tyrode’s solution, and imaged using structured light microscopy (Materials and Methods). Fig. 4A shows an illuminated plane (~ 700 nm thick) that is two microns above the focal plane of the coverslip. Perfusion of 10 mM lactate (Fig. 4B) resulted in a rapid loss of fluorescence that quickly reversed and then slowly approached steady state, indicative of lactic acid entering the cell. Washout of extracellular lactate led to a rapid efflux of protons from the cell that diminished slowly over time. Lactate-induced proton transport was inhibited with the monocarboxylate transporter (MCT) inhibitor (AR-C155858: 50 nM), suggesting that cardiac MCT-1 was responsible for lactic acid transport into the cell. In contrast, proton efflux after lactate washout was likely an amalgam of both cardiac MCT-1 and Nhe1 (Na^+^/H^+^) activity (Johannsson et al., 1997; Petrecca et al., 1999). To isolate the sarcolemma and t-tubule signals, we analyzed concentric annuli by systematically eroding five pixels from the cardiomyocyte edge (Fig. 4C). Annulus 0 (A0) contained primarily signal from the sarcolemma whereas the signal from the next three annuli (A1 – A3) were increasingly enriched with t-tubules (Fig. 4D). Because the mean of WGA-pHRho labeling varied less than 1% between annuli (Fig. 4E), the absolute change of fluorescence corresponds to the ion concentration for each annulus and was compared (Fig. 4F). Starting at the sarcolemma (A0), the initial change in fluorescence upon lactate perfusion increased significantly (p < 0.05) when compared to the inner annulus until A2, where there was no significant difference between the A2 and A3 annuli. In contrast, the initial change in fluorescence upon lactate washout was not significant when comparing adjacent annuli, but significant differences in initial fluorescence were measured between non-neighboring annuli. Taking together, these results demonstrate that the initial MCT-1 activity results in significantly more proton depletion in the t-tubules compared to the sarcolemma.

**Figure 4.**
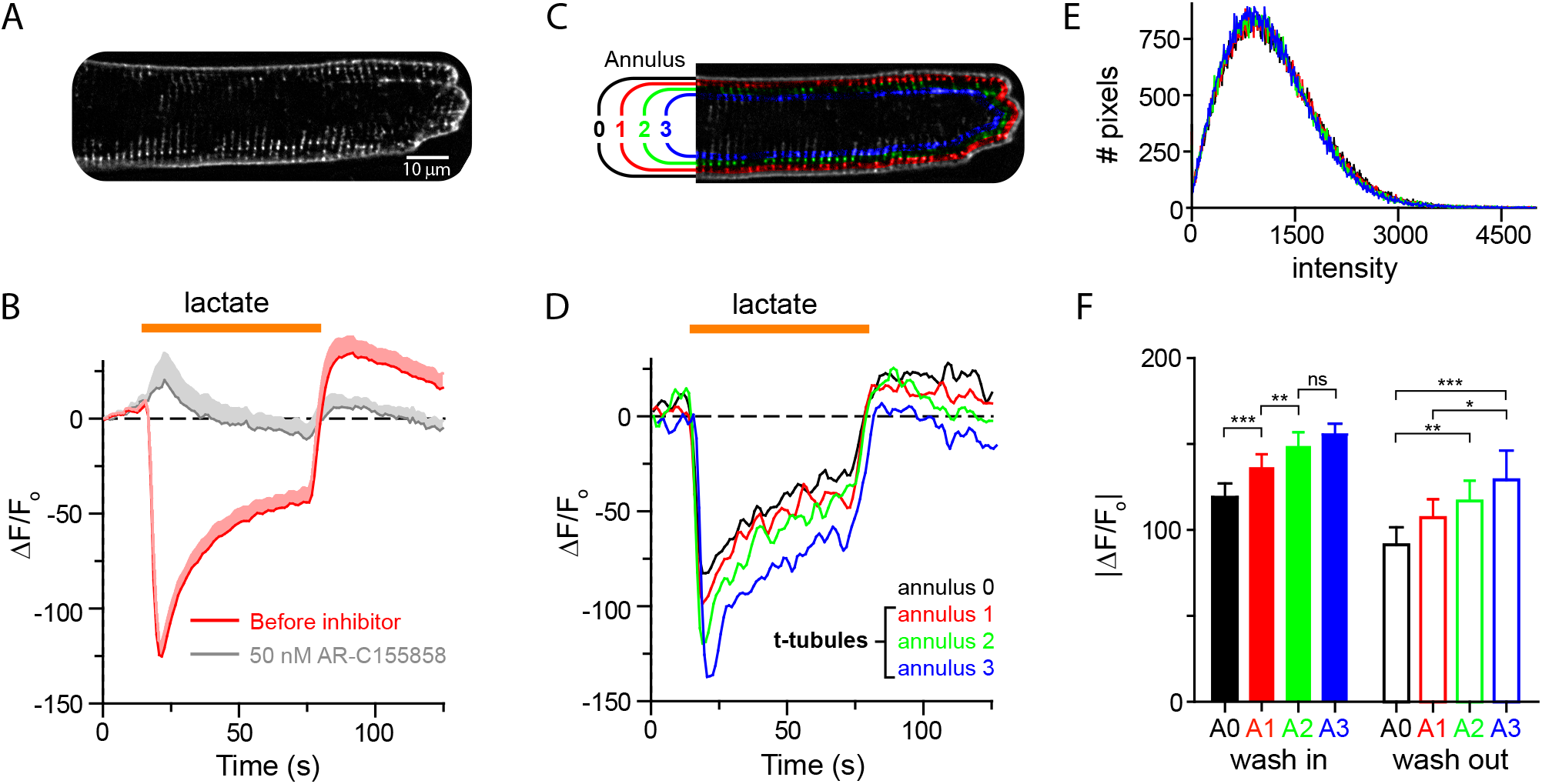
Structured light microscopy of lactate-stimulated proton transport in rat ventricular myocytes labeled with WGA-pHRho. (**A**) A ~ 750 nm illuminated *z*-plane of a WGA-pHRho labeled ventricular myocyte approximately two microns above the coverslip focal plane. (**B**) Time course of 10 mM lactate perfusion before (red) and after addition of 50 nM AR-C155858 (gray). Plotted data are mean + SEM (shading) for clarity (n = 6 cells); data were collected at 1Hz. (**C**) Psuedo-color image depicting the eroding pixel (5 px) analysis to generate the four annuli. (**D**) Changes of ΔF/F_0_ before, during, and after perfusion of 10 mM lactate at each annulus (A0-A3). (**E**) Pixel intensity histogram of the four annuli (area normalized) for cell shown in **A**. (**F**) Average maximal |ΔF/F_0_| upon wash in (solid bars) and wash out (open bars) of 10 mM lactate for nine cells. 1-way ANOVA (Bonferroni’s Multiple Comparison Test) was used to determine significance: *p < 0.05; **p < 0.01; ***p < 0.001; ns: not significant.

We next determined whether WGA-pHRho could detect proton efflux from primary neuron-astrocyte co-cultures. Co-cultures (6 – 10 d) obtained from mice cortex were labeled with WGA-pHRho in free glucose media, rinsed with free glucose media, and imaged by means of laser scanning microscopy. When co-cultured, neurons and astrocytes organize such that astrocytes occupy the bottom of the chamber, while most of the neurons grow on top. Therefore, two optical planes were imaged to capture both cell types. Figure 5A is a representative merged image of the neuronal and astrocyte planes labeled with WGA-pHRho and DAPI. To elicit lactic acid efflux, glucose-starved co-cultures were exposed to glucose (3 mM) while following the WGA-pHRho signal during acquisition. Merged neuronal and astrocyte planes in Figures 5B and 5C show the average change in fluorescence induced by the addition of glucose to the media (Fig. 5D). As expected, an increase in the substrate of the glycolytic chain is accompanied with an increase in the release of protons, an indirect readout for lactic acid release, presumably through monocarboxylate transporters (Dimmer et al., 2000). The overall effect of glucose addition was similar in both cell types; however, poisoning the mitochondrial respiratory chain with rotenone (100 μM) (Fig. 5D) led to clear differences in their response. Although lactic acid efflux from astrocytes was relatively unaffected by rotenone, the increase of lactic acid efflux from neurons is evident (Figs. 5E – 5H). These results are consistent with previous reports (Magistretti and Allaman, 2015) that mitochondrial oxidative capacity is greater in neurons than astrocytes in co-culture.

**Figure 5:**
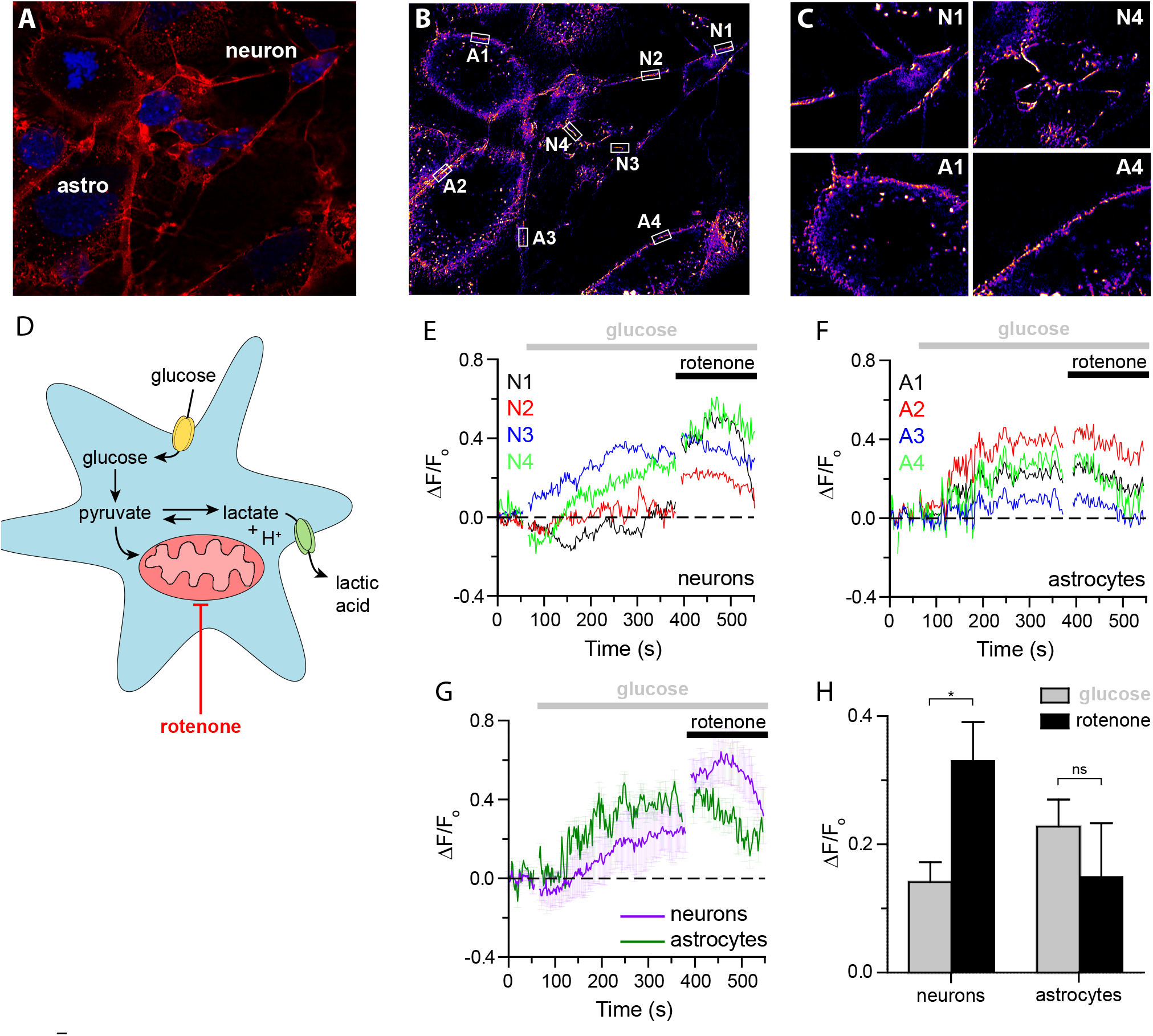
Proton efflux from primary neuron-astrocyte co-cultures labeled with WGA-pHRho. (A) Merged Airyscan image of the neuronal and astrocyte planes labeled with WGA-pHRho (red) and DAPI (blue). (B) Pseudo-color image showing the averaged ΔF/F_0_ of the ROI in A after glucose addition. Red shades indicate more activity over time. Regions of interest are denoted by white boxes (N = neuron; A = astrocyte). Data were collected at 2Hz. (C) Magnified images of two astrocytes and neurons in B to show labeling coverage. (D) Cartoon depiction of the metabolic pathways expected to be affected by glucose starvation and rotenone mitochondrial poisoning. Time courses of the (E) neuronal and (F) astrocytic pHRho signal during glucose (3mM) and rotenone (100 μM) addition. (G) Averaged data from panels E and F (total number of cells in the plot: 4 neurons and 4 astrocytes from one single dish; this average is considered n=1). (F) Overall change in ΔF/F_0_ (n = 4 – 6 plates), comprising a total of 19 neurons and 21 astrocytes in total; bars are SEM.

## Discussion

We sought to circumvent the secretory pathway and exploit WGA binding to mammalian glycocalyces to develop a simple, non-genetic approach to fluorescently visualize extracellular proton accumulation and depletion from isolated cells. Despite binding to sialic acid and N-acetylglucosamine (Monsigny et al., 1980), the voltage gating of glycosylated and nonglycosylated ion channels was not substantially altered after WGA treatment (Fig. 2B). WGA labeling of the cell surface was extremely uniform with only a 1% difference in labeling intensity (Fig. 4E) between annuli, enabling the change in fluorescence (ΔF/F_0_) to be converted into a pH change (Fig. 3). Because the buffer capacity must be kept extremely low to observe a robust signal, using this approach to measure the extracellular pH adjacent to the cell surface is not recommended. Compared to our previously published small molecule approach (Zhang et al., 2016), labeling with WGA conjugates does not require the cells to metabolize the unnatural sugar, which enabled facile labeling of primary cardiomyocytes and neuron-astrocyte cocultures. In addition, we initially observed very little intracellular labeling with WGA conjugates because the protein conjugates are not membrane permeant. However, both approaches target cell surface glycoproteins, so the fluorescent pH sensors are internalized over time, leading to unaesthetic bright puncta because WGA-pHRho is more fluorescent in acidic compartments. Although changes in fluorescence can be observed > 3 h after labeling (Fig. 2E), rapid cell surface labeling and subsequent imaging is the major advantage of using WGA conjugates to detect extracellular proton accumulation and depletion over the landscape of a cell.

To demonstrate the utility of WGA conjugates for detecting spatiotemporal differences in extracellular proton accumulation and depletion, we labeled primary ventricular cardiomyocytes and neuron-astrocyte co-cultures with WGA-pHRho and imaged at several *z*-planes. Using structured illumination at single *z*-plane, we observed significantly more lactic acid depletion in annuli enriched with t-tubules. Although the initial influx of lactic acid can be attributed to MCT-1, the larger pH change observed in the t-tubules could be due to either more MCT-1 transporters localized to the t-tubules and/or the restricted exchange with bulk solution due to the architecture and membrane folds of the t-tubule network. The pixel intensity histogram (Fig. 4E) shows uniform labeling of the annuli with WGA-pHRho; however, it is unclear whether WGA-pHRho labels the entire cardiac tubular network. A lipid-conjugated pHRho or trapping a small molecule, membrane impermeant, fluorescent pH sensor in the extracellular t-tubule network (Launikonis et al., 2018) may be better suited to investigate the narrowest cardiac t-tubules as well as the skeletal t-tubular system. For neuron-astrocyte co-cultures, WGA-pHRho and imaging on two alternating planes revealed differences between neuronal and astrocytic proton fluxes and mitochondrial oxidative capacity. In both experimental paradigms, we used WGA-pHRho to detect the proton that is co-transported with lactate, which is prone to contamination by other proton-coupled (and bicarbonate) transporters in primary cells that are attempting to regulate intracellular pH. WGA conjugates with small molecule (Pal et al., 2009) or proteinbased (San Martin et al., 2013) fluorescent lactate sensors would allow more specific detection of extracellular lactic acid accumulation and depletion in a well-buffered external solution.

In summary, WGA-pHRho is easy to make and it quickly and specifically labels the glycocalyx of primary cells in culture. WGA-pHRho is ideal for detecting proton accumulation and its photostability makes it compatible for use with several imaging modalities. As a non-genetically encoded approach, it is amenable for usage with human cells and humans as WGA-fluorescein has been used to visualize the ocular glycocalyx of soft contact lens wearers (Fukui et al., 2016). The straightforward labeling and purification procedures outlined herein will allow for the synthesis of WGA pH sensitive conjugates with various pKa values and excitation and emission maxima. Moreover, the approach is readily generalizable to other small molecule ion-sensitive fluorophores that possess pendant carboxylic handles that can be conjugated to WGA.

## Author contributions

LZ synthesized pHRho and developed the WGA derivatization procedure. MZ performed the CHO and cardiomyocyte experiments. Patch clamp fluorometry setup was built by KB; structured illumination data were acquired on TESM, analyzed by KF, and was built and is maintained by KF and KB. MAC designed the glucose/rotenone experiments and prepared the astrocyte neuron co-cultures. SB labeled, imaged, and analyzed the co-culture data. WRK designed the experiments, created the figures, and wrote the manuscript.

All authors have approved the final version of the manuscript.

## Acknowledgements

The authors declare that they have no competing financial interests. This work was supported by a grant to WRK from the National Institutes of Health (GM-070650) and by the US/Chile Fulbright Scholar Program (WRK).

## Supplemental Figures

**Figure S1, related to Figure 2.**
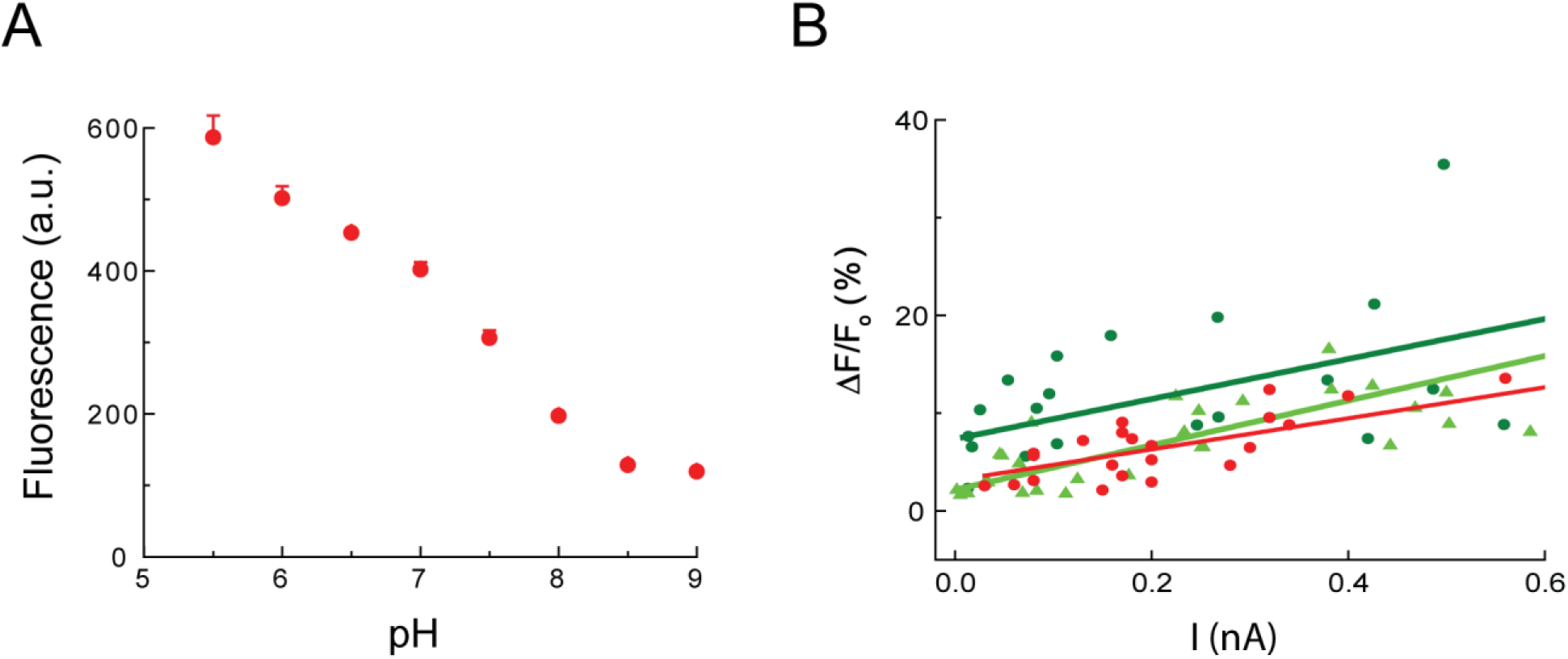
Characterization of pH sensitive WGA conjugates. **(A)** Fluorescence-pH plot of WGA-pHRho in solution; excitation wavelength: 550 nm. **(B)** ΔF/F_0_-I plot of WGA-pHRho (red), homemade WGA-fluorescein (dark green) and WGA-fluorescein from Vector Laboratories (light green).

**Figure S2, related to Figure 2.**
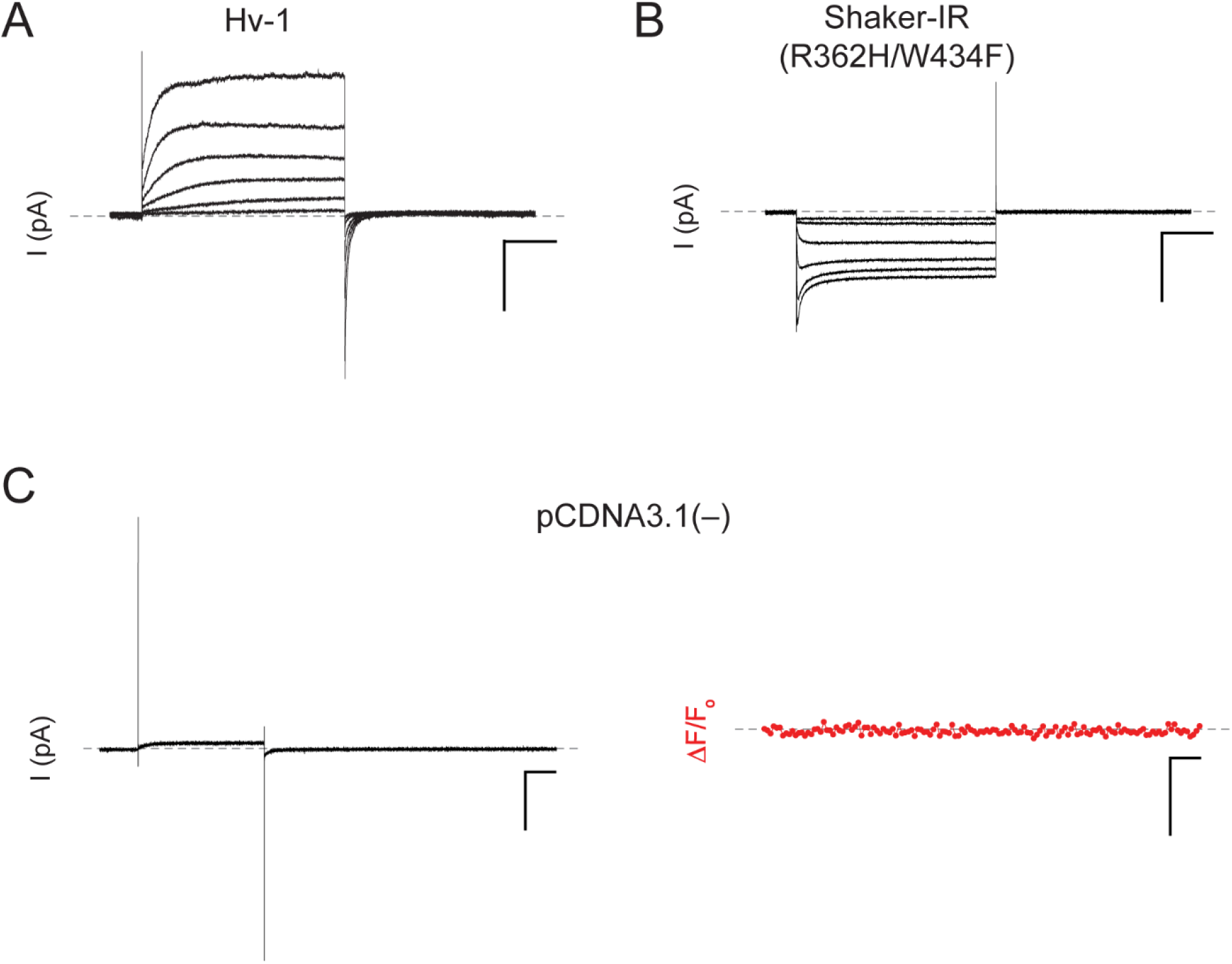
Current-voltage relationships for Hv-1 and Shaker-IR (R362H/W434F) expressed in CHO cells. **(A)** The cell was held at – 80 mV and currents were elicited from 4-s command voltages from 0 to 100 mV in 20-mV increments; pH_o_/pH_i_ = 7.5/6.0 (0.1 mM HEPES). **(B)** The cell was held at + 30 mV and the currents were elicited from 4-s command voltages from – 120 to – 20 mV in 20-mV increments; pH_o_/pH_i_ = 6.0/7.5 (0.1 mM HEPES). Scale bars in **A** and **B** are 250 pA and 1 s. **(C)** Voltage-clamp fluorometry of a CHO cell transfected with pCDNA3.1(–) control. The cell was held at – 80 mV and depolarized to 100 mV for 4 s. Scale bars represent 100 pA, 5% and 1 s; pH_o_/pH_i_ = 7.5/6.0 (0.1 mM HEPES).

## Supplementary movie legends

**Supplementary movie 1, related to Figures 2C and 3C.** A ΔF/F_0_ movie of a representative CHO cell expressing Hv-1. The cell was held at – 80 mV and depolarized at 100 mV for 4 s. F_0_ is the fluorescent signal of the first frame; the frame rate is 10 fps. The patch clamped cell becomes the brightest cell in the center of the frame upon depolarization. Under the low buffer capacity conditions (0.1 mM HEPES), the fluorescence of the WGA-pHRho labeled neighboring cells record the proton wavefront emanating and dissipating from the patch clamped cell.

**Supplementary movie 2, related to Figure 2C and 3C.** Raw fluorescent images of a representative CHO cell expressing Shaker-IR (R362H/W434F) labeled with WGA-pHRho. The cell was held at 30 mV and hyperpolarized to – 120 mV for 4 s. The patch clamped cell is denoted with an arrow; the frame rate is 10 fps.

